# Structural study of UFL1-UFC1 interaction uncovers the importance of UFL1 N-terminal helix for ufmylation

**DOI:** 10.1101/2022.09.15.508077

**Authors:** Sayanika Banerjee, Julia K Varga, Manoj Kumar, Guy Zoltsman, Michail N Isupov, Rina Rosenzweig, Ora Schueler-Furman, Reuven Wiener

**Affiliations:** Department of Biochemistry and Molecular Biology, The Institute for Medical Research Israel-Canada, Hebrew University-Hadassah Medical School, Jerusalem 91120, Israel; Department of Microbiology and Molecular Genetics, The Institute for Medical Research Israel-Canada, Hebrew University-Hadassah Medical School, Jerusalem 91120, Israel; Department of Chemical and Structural Biology, Weizmann Institute of Sciences, Rehovot, Israel; The Henry Wellcome Building for Biocatalysis, Biosciences, University of Exeter, Stocker Road, Exeter, EX4 4QD, United Kingdom

## Abstract

Ufmylation, a protein modification by Ubiquitin-like (UBL) protein UFM1, plays a crucial role in several cellular processes including DNA damage response, protein translation and ER homeostasis. To date, little is known how the enzymes responsible for this modification coordinate their action. Here we have studied the details of UFL1 (E3) activity, its binding to UFC1 (E2), and its relation to UBA5 (E1), using a combination of structural modeling with Alphafold2, X-ray crystallography, NMR, and *in vitro* biochemical activity assays. Guided by an Alphafold2 model, we generated an active UFL1 fusion construct that includes its cofactor DDRGK1, and solved the first crystal structure of this critical interaction. This fusion construct also unveiled the importance of the N-terminal helix of UFL1 for its binding to UFC1, which was validated by ITC and NMR experiments. Importantly, the binding site suggested by our structural model of the UFL1-UFC1 interaction reveals a conserved interface, and suggests a competition for binding to UFC1 between UFL1 and UBA5, which we reconfirmed by NMR. Altogether, our study reveals a novel, terminal helix-mediated regulatory mechanism which coordinates the cascade of E1-E2-E3 mediated transfer of UFM1 to its substrate, and provides new leads to target this important modification.

**Significance statement:** Ufmylation is an important post-translational modification, but little is known about the mechanistic details of its machinery, and in particular how the UFM1 E3 ligase (UFL1) binds and functions together with the E2 conjugating enzyme (UFC1). We combined AlphaFold2 modeling, X-ray crystallography, NMR and biochemical experiments to reveal crucial elements that govern UFL1 activity and ufmylation. We discover a crucial role for the UFL1 N-terminal helix in binding to UFC1 and productive ufmylation. This helix competes with the E1 (UBA5) C-terminal helix for binding to UFC1. Altogether, our findings uncover a new, helix-mediated regulatory mechanism in ufmylation.

## Introduction

Protein modifications by UFM1 (ufmylation) play a role in many cellular processes, including DNA damage repair, the anti-viral and the unfolded protein responses (1). A three-enzyme cascade involving the E1-UBA5, the E2-UFC1 and the E3-UFL1 is responsible for the attachment of UFM1 to target proteins. Initially, UBA5 activates UFM1 in an ATP-dependent process. Then, UFM1 is transferred from UBA5 to the active site cysteine of UFC1, forming a thioester bond. Finally, with the help of UFL1, UFM1 is transferred from UFC1 to the target protein (2, 3).

Surprisingly, UFL1 lacks structural elements that are common to other E3 ligase enzymes, namely a RING domain, a HECT-type catalytic domain or an RBR structure (4–6). In addition, while some atypical E3 enzymes possess a motif required for interaction with their ubiquitin-like protein (7, 8), whether UFL1 has a UFM1-interacting motif is uncertain. Therefore, it remains to be determined whether UFL1 functions in a novel mechanism that does not exist in other E3 ligases. It was shown previously that ufmylation by UFL1 of the nuclear receptor coactivator, ASC1, requires DDRGK1 (also known as UFBP1) (9). Moreover, a recent model of the interaction between UFL1 and DDRGK1 generated by AlphaFold2 has revealed structural complementation between the two proteins (10). Besides binding to DDRGK1, UFL1 interacts with LZAP (also known as the adapter protein CDK5RAP3), forming a ternary complex (11). The latter has been suggested to possess a motif allowing UFM1 binding (12). Currently, structural data on UFL1 is still missing, and it is still unclear how UFL1 binds to UFC1 to promote ufmylation.

Deep learning of modeling of protein structures is revolutionizing the field of structural biology, spearheaded by AlphaFold2 developed by DeepMind (13–15). As models are either available, or can be generated within a short time on platforms such as ColabFold (16), they will accelerate studies that previously depended on the expression, purification, and crystallization of one or more proteins, a process that could take years, if successful at all. Structural models provide guidelines for the generation of stable proteins, by identifying disordered regions that hamper protein expression and could be truncated for improved expression. Beyond the study of protein monomers, the structures of many protein complexes can now be modeled (17–19), including interactions mediated by short motifs (20, 21). It is now possible to also study the interaction of regions that adopt a stable structure only upon interaction, as for example the interaction of motifs located within disordered regions with their partners, as well as complementation of a full domain by two proteins.

With these tools in hand, we set out to study the ufmylation system and reveal yet unsolved challenges in our understanding of this complex regulatory pathway. We report here on two major advances: 1) The design of a functional fusion protein, composed of truncated parts of the UFL1-DDRGK1 complex. This construct allows to significantly simplify the study of UFL1 activity, and importantly, has enabled us to solve, for the first time, a crystal structure of UFL1 bound to DDRGK1; 2) The definition of the critical role of the UFL1 N-terminal helix in UFC1 binding and ufmylation. Our model suggests that E2 UFC1 uses the same site to bind both E1 UBA5 as well as E3 UFL1 helices, which we confirm by NMR studies. Of note, this helix-mediated interaction was revealed in a model generated without any prior information, highlighting the contribution of AlphaFold2 to the revelation of new interaction details and regulation.

### Results

### AlphaFold2-assisted engineering of an active UFM1 E3-ligase

UFL1 has been suggested to be only active in the presence of DDRGK1(22). In line with a parallel recent study (10), our starting point was a model of the UFL1-DDRGK1 interaction (see **Figure 1a** and **Supplementary Figure 1)** that we generated using AlphaFold2 (see Methods). The model shows the crucial contribution of DDRGK1 to complement the first winged helix domain repeat of UFL1 (residues 27-58) and explains why neither DDRGK1 nor UFL1 are folded when expressed alone. This model, as well as additional information about the importance of different regions (3), assisted us in the design of a fusion construct encompassing DDRGK1-UFL1 (**Figure 1b**), in which we removed the N-terminal region of DDRGK1 and the C-terminal region of UFL1 (*i.e.,* DDRGK1:207-314 - UFL1:1-200). Furthermore, we noted the predicted low confidence of the N-terminal helix of UFL1 (average AlphaFold2 pLDDT < 70) (**Figure 1c**), and therefore generated a second construct in which we also truncated this N-terminal UFL1 helix, DDRGK1-UFL1ΔN (DDRGK1:207-305 - UFL1:27-200). To test the AlphaFold2-based design of the above fusion proteins comprising UFL1-DDRGK1, we purified the fusion proteins (**Supplementary Figure 2**) and tested their elution profile using gel filtration, demonstrating that the fusion proteins do not elute as soluble aggregates (**Figure 1d**). Overall, our results imply that, similar to the co-expression of UFL1 with DDRGK1 that allows purification of a soluble UFL1-DDRGK1 complex (10), the fusion protein is also soluble.

**Figure 1:**
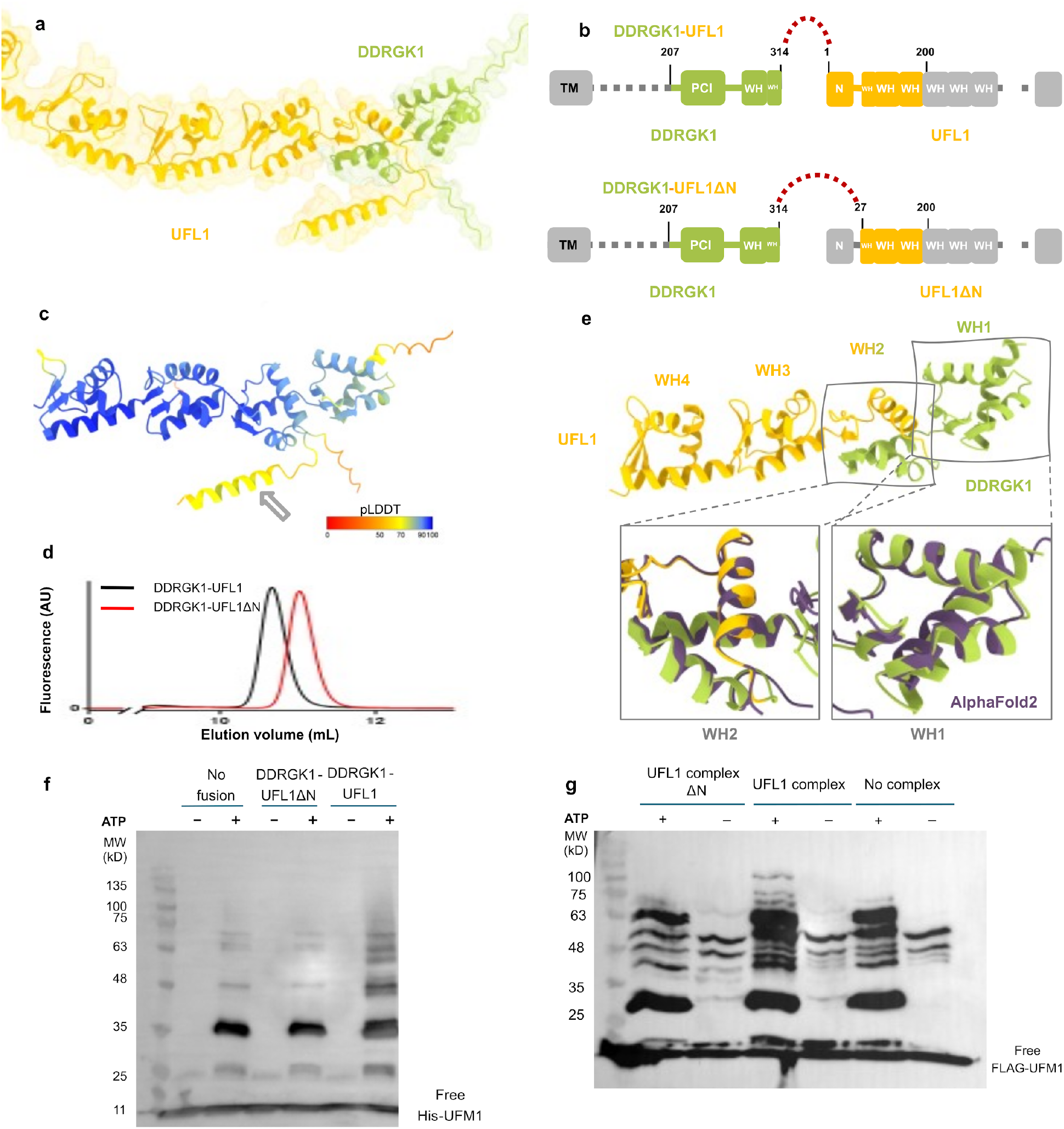
AlphaFold2-assisted generation of an active fusion protein for ufmylation. **a** AlphaFold2 structural model of the DDRGK1-UFL1 complex (similar to Peters *et al. (10)*; DDRGK1 colored in green; UFL1 colored in yellow). **b** Details of the proteins and the designed fusion constructs. DDRGK1 was connected to UFL1, removing the N-terminal region of DDRGK1 and the C-terminal region of UFL1. In a second construct we also removed the N-terminal helix of UFL1 (DDRGK1-UFL1ΔN), due to its flexibility suggested by AlphaFold2 (see Text). Regions removed from the parent proteins are shown in gray (WH: winged helix domains, PCI: proteasome-COP9-initiation factor 3 domain). **c** Model of the DDRGK1-UFL1 complex, colored according to pLDDT, highlighting the low confidence in the structure of the N-terminal helix (gray arrow). **d** Gel filtration elution profiles of fusion proteins. **e**. Crystal structure of the DDRGK1-UFL1ΔN fusion. The four winged helix (WH) domains are indicated and numbered. The blow-ups are superimposed onto the AlphaFold2 model of the indicated WH domains (purple). **f-g** In vitro ufmylation assays. Western blots show that ufmylation activity depends on the presence of the N-terminal helix of UFL1 in **f** the fusion protein, and **g** the ternary complex of UFL1, DDRGK1 and LZAP. See **Supplementary Figure 3** for protein loading controls.

To date, structural data on DDRGK1-UFL1 complexes are based on AlphaFold2 models (10). This motivated us to exploit our fusion proteins for determination of their crystal structure. We successfully solved the crystal structure of DDRGK1-UFL1ΔN to 3.1 Å resolution (**Supplementary Table 1**). The structure reveals four repeats of winged helix (WH) domains: the first is contributed by DDRGK1, the second is formed partially by DDRGK1 and partially by UFL1, while the last two are from UFL1 (**Figure 1e**). This structure is very similar (backbone RMSD = 1.4 Å) to our corresponding AlphaFold2 model, suggested also by Peter *et al.* (10). Most of the structural differences are concentrated in the first WH domain (residues 1-65 in the fusion; backbone RMSD = 2.4 Å). Interestingly, less structural differences are observed in the combined WH domain (residues 66-130 in the fusion; backbone RMSD = 2.1 Å), although both of its parts are connected in the fusion protein (**Figure 1e**). In the crystal structure the last 18 amino acids are flexible and are not detected in the electron density map. Overall, our crystal structure suggests that the fusion protein maintains the overall architecture of the UFL1-DDRGK1 complex as observed in the AlphaFold2 model.

**Table 1:**
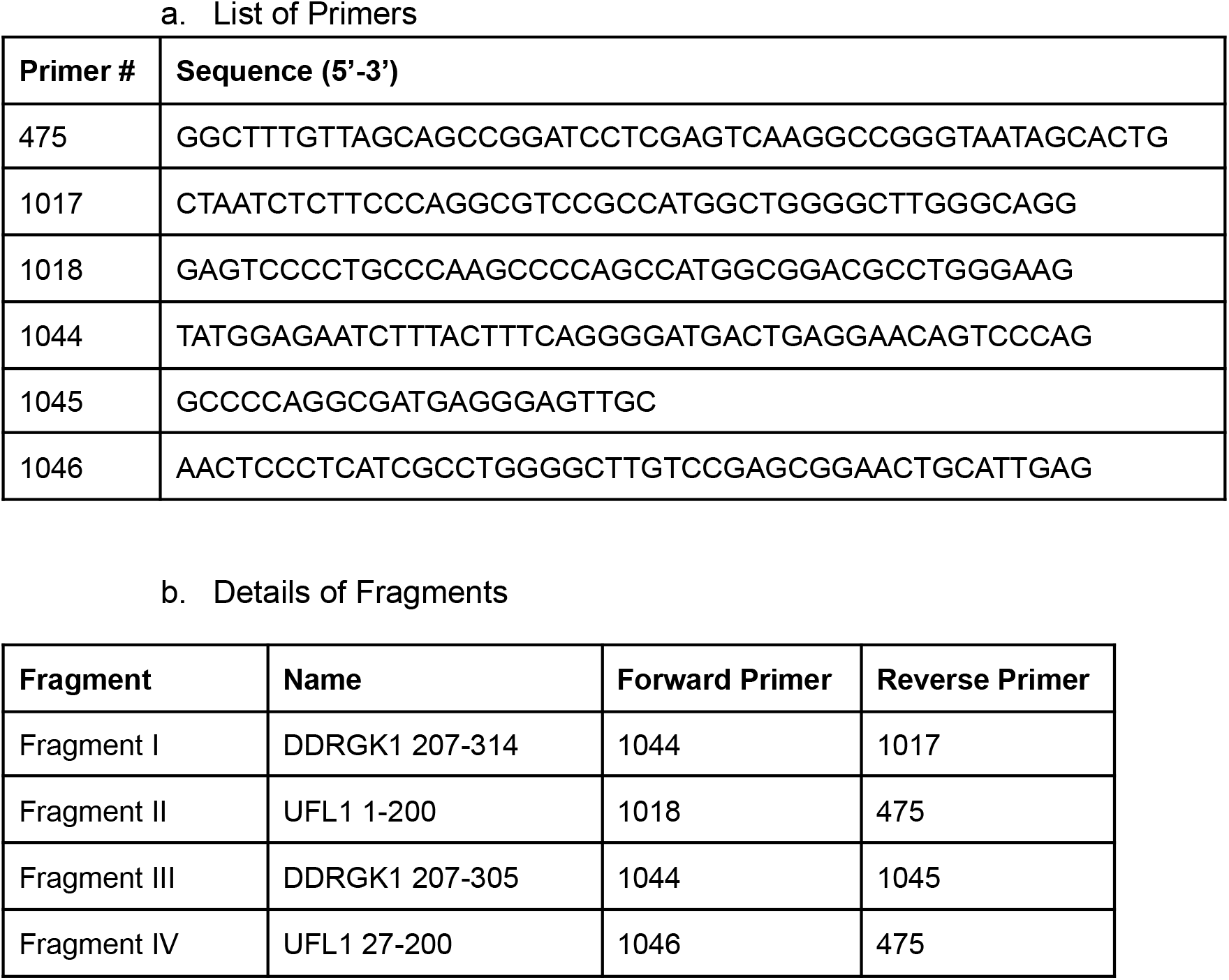
Primers and Fragments used/generated in this study

With the above fusion proteins in hand, we tested their functionality as E3 ligases. To that end we incubated pure UBA5, UFC1 with and without fusion proteins and analyzed the ufmylation pattern (**Figure 1f**). Reassuringly, addition of our full fusion construct (DDRGK1-UFL1) resulted in changes in the ufmylation pattern, as did purified ternary complex (involving UFL1, DDRGK1, and LZAP **Figure 1g**). To our surprise, however, the fusion protein lacking the UFL1 N-terminus (DDRGK1-UFL1ΔN) did not show any such changes (**Figure 1f**), and neither did the ternary complex lacking the N-terminal helix (**Figure 1g**), suggesting that the UFL1 N-terminus is essential for E3 ligase activity.

### A structural model of the UFC1-UFL1 interaction reveals a critical role of the helix in the N-terminal tail of UFL1

Why is the UFL1 N-terminal helix critical to its ligase activity? Motivated by the contribution of the AlphaFold2 model to the successful design of a fusion protein with E3 ligase activity, we decided to model the binding of UFC1 to the UFL1-DDRGK1 complex (**Figure 2a,b**). This model explains the dominant contribution of the N-terminal helix to this interaction, and identifies the residues in UFC1 that are crucial for its binding (Y36, I40, R55, **Supplementary Table 2**). The importance of the interface hotspot residues in the UFL1 helix is emphasized by their high degree of evolutionary conservation (W5, I8, L11 and F15; **Figure 2c)**. As mentioned above, this N-terminal region was modeled as a helix by AlphaFold2, albeit with low confidence in the apo structure (**Figure 2d**, see also **Figure 1c**) which indicated that it might be disordered and only fold into a helix upon binding. Our ITC binding experiments confirmed that only the fusion protein possessing the UFL1 N-terminal helix binds to UFC1 (Kd = 2.4μM; **Figure 2e, Supplementary Table 3**). We reconfirmed this also by pulldown experiments (**Supplementary Figure 4**). Of note, we previously measured similar binding affinity for the interaction of UFC1 with UBA5 (0.94 μM for full UBA5) and 5.5 μM for the C-terminal region of UBA5 (residues 347-389)(23, 24).

**Figure 2:**
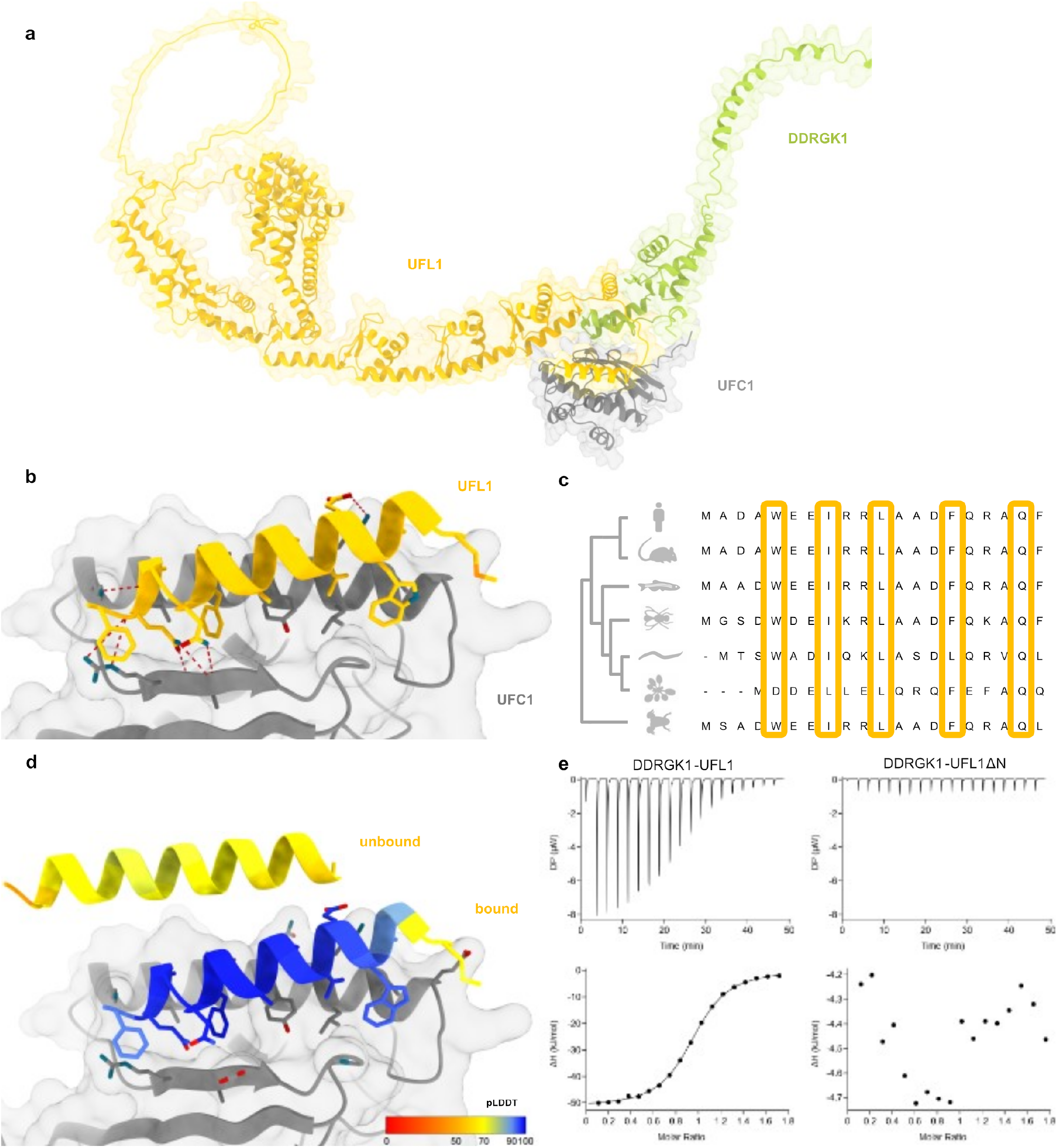
The N-terminal helix of UFL1 is crucial for binding of UFC1 and activity. **a** Overall view of the UCF1-DDRGK1-UFL1 ternary complex. **b** Details of the interaction: UFL1 N-terminal helix bound to UFC1. **c** Analysis of conservation of UFL1 N-terminal helix shows evolutionary conservation of the residues predicted to be involved in binding. **d** The N-terminal helix (colored according to pLDDT) is modeled with high confidence when bound to UFC1 (in gray), in contrast to the low confidence for this helix in the unbound structure (see also **Figure 1c**). **e** ITC experiments of UFC1 binding to DDRGK1-UFL1 (left panel) and DDRGK1-UFL1ΔN (right panel). The top graph represents raw data of heat flow versus time. The area under the peaks of the upper panel was integrated and plotted as kJ per mole of UFC1 as a function of binding stoichiometry in the bottom panel. Thermodynamic parameters are summarized in Supplementary **Table 3.** In panels (**b)** and **(d**) AMBER relaxation was performed after AlphaFold2 structure prediction, to optimize side-chain orientations.

To support our model of the interaction of UFL1 N-terminus with UFC1, we used NMR spectroscopy to define the changes in UFC1 chemical shifts upon binding to DDRGK1-UFL1 or DDRGK1-UFL1ΔN. To that end, we exploited the reported assigned (^1^H, ^15^N)-HSQC NMR spectra for UFC1(25). As expected from our activity and ITC experiments **(Figures 1f and 2e**), the addition of DDRGK1-UFL1, but not DDRGK1-UFL1ΔN, to ^15^N-labeled UFC1 caused strong attenuations (**Figure 3a,b**). In line with the AlphaFold2 model, these attenuations include residues from UFC1 **α**-helix I (residues 26-48) and from **β**-strand I (residues 54-58), that directly interact with the UFL1 N-terminus (**Figure 3c**). Interestingly, besides the above residues, the N-terminal half of UFC1 **α**-helix II (residues 135-145) also showed chemical perturbations. This region, according to our AlphaFold2 model, is located on the other side and not directly involved in UFL1 N-terminus binding, raising the possibility that these residues are allosterically regulated by binding of the UFL1 N-terminus to UFC1.

**Figure 3:**
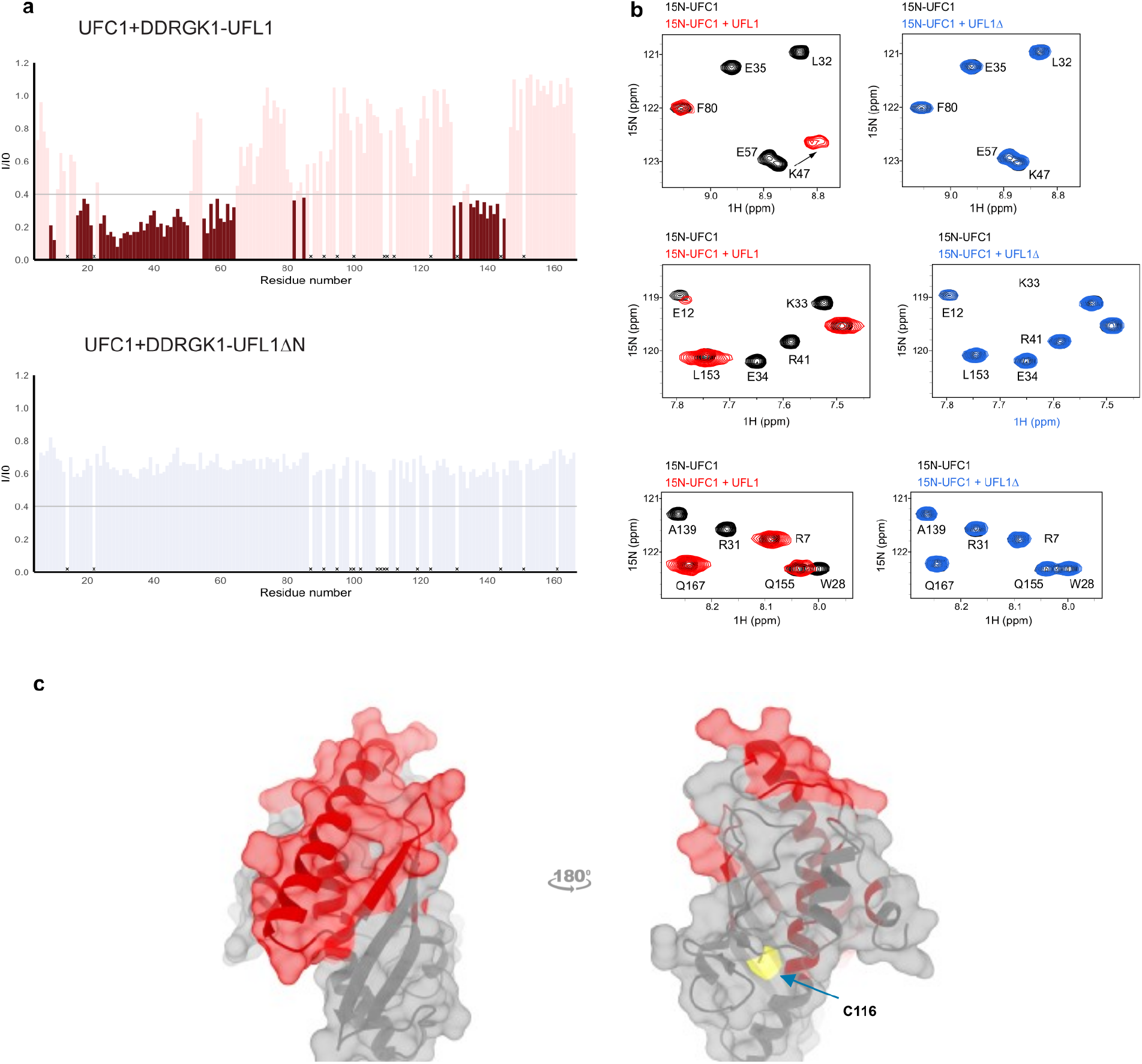
Characterization of UFC1 binding to DDRGK1-UFL1 using NMR. **a** Intensity changes of UFC1 residue peaks upon addition of 1.5-fold excess (300 μM) of DDRGK1-UFL1 (maroon) or 2-fold excess (400 μM) DDRGK1-UFL1ΔN (blue). Dark colors indicate shifted residues (I/I_0_ <= 0.4), light colors indicate unshifted residues (I/I_0_ > 0.4), ‘x’ denotes residues without assigned values. Removal of the N-terminal UFL1 regions significantly impairs UFC1-DDRGK1-UFL1 complex formation. **b** Selected regions of the ^1^H–^15^N HSQC spectrum: 0.2 mM UFC1 alone (black) and in the presence of two-fold excess of DDRGK1-UFL1 (red) or DDRGK1-UFL1ΔN (blue). **c** structure of UFC1 [PDB ID: 7NW1 (23)] with residues displaying significant intensity changes (I/I_O_ <= 0.4) upon addition of DDRGK1-UFL1 colored in red.

According to our model and NMR data (**Figures 2b** and **3**), UFC1 binds the N-terminal helix of UFL1 using the same binding pocket with which it also binds to the C-terminal helix of UBA5 (as shown by a crystal structure solved previously by us (23), **Figure 4a**). This suggests a competition between UFL1 and UBA5 for binding to UFC1. To verify this, we performed an NMR based competition experiment between ^15^N-labeled UBA5 C-terminus (UBA5 347-404) bound to UFC1, and UFC1+UFL1 (**Figure 4b**). As expected, addition of UFC1 to ^15^N UBA5 C-terminus caused large changes to the UBA5 ^1^H–^15^N HSQC spectrum, in the form of chemical shift perturbations (**Figure 4b, center panel**). Addition of UFL1 to UBA5-UFC1 complexes, however, caused shifting of NMR cross-peaks in UBA5 to their unbound position, conforming that UFL1 and UBA5 bind to the same surface of UFC1 (**Figure 4b, right panel**). To summarize, these results demonstrate that both E1 UBA5 and E3 UFL1 use the same pocket on E2 UFC1 for binding, revealing important details of the regulation of ufmylation.

**Figure 4.**
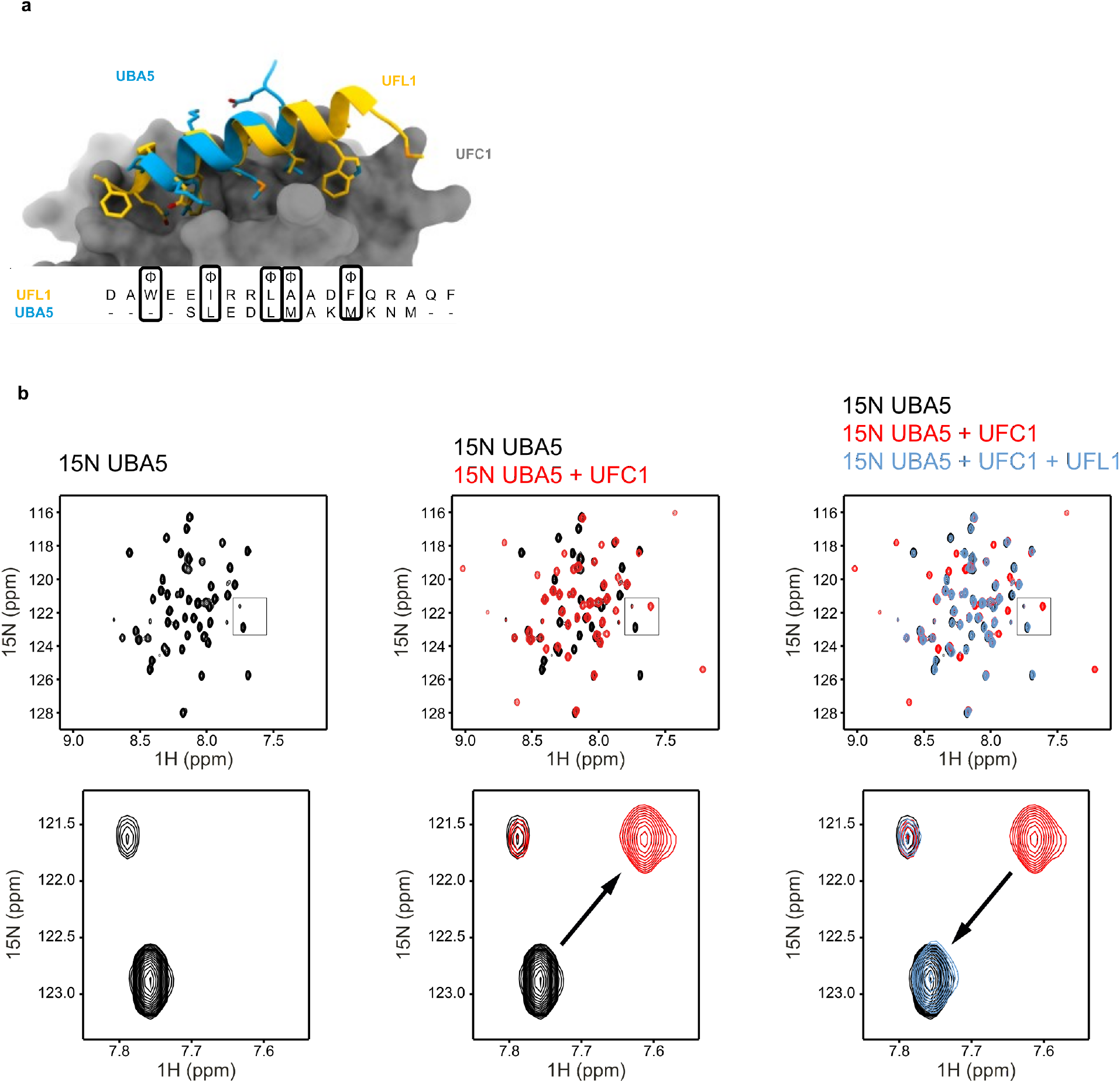
UBA5 and UFL1 compete for binding to UFC1. **a** Superposition of the model of the UFL1 N-terminal helix-UFC1 complex onto the solved structure of the UBA5 N-terminal helix-UFC1 complex (PDB ID: 7NW1) suggests that both bind to the same binding site (as has been suggested also for ubiquitin E1, E3 enzymes (26))**. b** NMR ^1^H-^15^N HSQC spectra revealing competitive binding: Left: ^1^H–^15^N HSQC spectrum of ^15^N UBA5 alone (black); Center: Overlay of spectrum after addition of UFC1 (red); Right: Overlay of spectrum after addition of both UFC1 and DDRGK1-UFL1 (blue), showing a return to the unbound spectra. Upper panel: Full spectrum; Lower panel: Inset focusing on a specific peak, showing its shift in the presence of UFC1, which is abolished after addition of competing UFL1.

## Discussion

Transient interactions between E1, E2 and E3 are indispensable for protein modifications by Ub and UBLs, therefore understanding their binding mechanisms is of high interest. Here we show that the UFL1 N-terminal helix that resides outside the winged helix domains is responsible for the binding to UFC1 (**Figures 1 and 2**). This helix binds UFC1 on the side opposite to the active site cysteine at position 116 (**Figure 3**), leaving the latter exposed for aminolysis. Intriguingly, this mode of binding overlaps with the binding site of UBA5 C-terminal helix on UFC1, suggesting competition between UBA5 and UFL1 on UFC1 (**Figure 4**). This competition allows UFC1 to execute its dual role in the conjugation process. Initially, UFC1 has to bind the charged UBA5 and participate in the trans-thiolation reaction where UFM1 is transferred to UFC1 active site cysteine. Then, UFL1 outcompetes UBA5, leaving UFC1 active site cysteine exposed and amenable to substrate’s nucleophilic attack. To date detailed understanding of how UFL1 outcompetes UBA5 needs further investigation. However, one possibility is that UFC1 affinity to UBA5 decreases when the latter is no longer charged, thereby allowing UFL1 to displace UBA5 and bind UFC1.

It is the context that will change the relative binding affinity to E2 by E1 and E3. While affinities measured in this and previous studies all lie in a similar range, the affinity measured for binding of UFL1-DDRGK1 to UFC1 (Kd = 2.4μM) is a little stronger than the corresponding affinity of binding of UBA5 C-terminus to UFC1 (Kd = 5.5μM). This is in agreement with a previous simulation, in which we ran AlphaFold2 on a sequence that contains both peptides, assuming that the stronger binding peptide will bind to the receptor binding site [an approach suggested by Chang *et al.* (27)]. In that experiment, the UFL1 helix invariably outcompeted the UBA5 helix in the binding site, and was modeled with significantly higher confidence (**Supplementary Figure 5**). This is anticipated since UFL1 binds as a longer helix (20 residues *vs.* 12 residues of UBA5) and generates a larger hydrophobic interaction surface. Within the setting of full proteins, charged or uncharged, different interactions will be favored along different steps in the ufmylation process.

Mechanistic studies on UFL1 as the E3 ligase of UFM1 have remained elusive until not long ago. While working with the intact UFL1 complex would be ideal, this is not always feasible for detailed structural research. The design of E3 constructs that are suitable for such study is therefore of high interest. However, making these constructs turns out to be highly challenging once the E3 of interest needs other partners for its folding and function as in the case of UFL1. Using AlphaFold2 we have generated a structural model for the E3 UFL1 together with its partner DDRGK1, and used it as the basis for the design of fusion proteins (**Figure 1**). These fusion proteins are highly expressed in *E. coli* and easily purified, bypassing the intrinsic challenges of working with the UFL1 complex. Importantly, they provided the first crystal structure of the UFL1-DDRGK1 complex (**Figure 1e**). Finally, they retain the E3 activity, and are thus suitable for a mechanistic study focusing on E2 - E3 interaction and function.

While we have previously demonstrated the ability of AlphaFold2 to model short motif-mediated interactions at high confidence (20), it was still surprising that AlphaFold2 could identify the binding region on UFL1 from the full-length protein. The resulting model provides the atomic details of the UFL1 helix - UFC1 interaction (**Figure 2**) and allows to identify the interface hotspot residues that are critical for this interaction (**Supplementary Table 2**). As we proceed to the study of additional components involved in the regulation of ufmylation, models generated by AlphaFold2 will continue to guide us and reveal additional details of regulation.

## Material and methods

### AlphaFold2 predictions

In general, structural models of individual proteins and complexes were generated using ColabFold(16). Due to the size of the UFL1-UFC1-DDRGK1 complex, all AlphaFold2-Multimer-v2.2 predictions on this complex were performed locally, using the LocalColabFold installation (downloaded on 17/07/2022 from https://github.com/YoshitakaMo/localcolabfold). Unless indicated otherwise, the predictions were run with all 5 models and default seed, default multiple sequence alignment generation using the MMSeqs2 server and with 3 recycles, without linkers between the monomers. The “computational competition assay” was run as suggested by Chang *et al. (27)*, providing both competing peptides in a single prediction run, provided before and after the receptor UFC1 sequence. All structure visualizations were created with ChimeraX v1.3 (28).

### Calculation of sequence conservation

Conservation of UFL1 was calculated with the ConSurf server(29), using default parameters. Alignment of the human and model animal sequences of UFL1 was performed using ClustalOmega (30) on the UniProt server (31).

### Computational alanine scanning

To estimate the contribution of different residues to the binding of UFL1 N-terminal helix to UFC1, we applied alanine scanning using the Robetta server (32). Residues with predicted effect of ΔΔG_binding_ > 1.0kcal/mol were retained as hotspot residues.

### Cloning

The fusion constructs DDRGK1_207-314_-UFL1_1-200_ (corresponding to DDRGK1-UFL1 in the main text) and DDRGK1_207-305_-UFL1_27-200_ (corresponding to DDRGK1-UFL1ΔN in the main text) (**Figure 1b**) were generated using Gibson assembly (Gibson assembly master mix, New England Biolabs) according to the manufacturer’s protocol. To generate the fragments by PCR, we used the Primers detailed in **Table Ia**, to generate fragments detailed in **Table Ib.** DDRGK1_207-314_-UFL1_1-200_ was cloned in pET15b by Gibson assembly of fragment I, II and linear pET15b. DDRGK1_207-305_-UFL1_27-200_ was cloned in pET15b by Gibson assembly of fragment III, IV and linear pET15b. All of the constructs were verified by DNA sequencing.

UFL1 1-794, UFL1 27-794 were cloned in pLVX with N-ter 3myc tag and C-ter strep tag in BamH1/ Xba1 site. LZAP with C ter His tag and DDRGK1 50-314 were cloned in pCDNA 3.4 by gibson assembly.

### Cell culture and transfections

293T cells were grown in Dulbecco’s modified Eagle’s medium supplemented with 10% fetal calf serum, and incubated at 37 °C in the presence of 5% CO_2_. 293T cells were grown to 70% confluence in T175 flasks and transiently co-transfected with 12 μg of 3myc_UFL1 1-794_Strep_pLVX/ 3myc_UFL1 27-794_Strep_pLVX, 24 μg of LZAP-His_pCDNA 3.4 and 24 μg of DDRGK1 50-314_ pCDNA 3.4 (total 60 μg of plasmid DNA) and 120 μl Transporter ™ 5 transfection reagent per flask. For full length UFL1 complex 40 x T175 were transfected and UFL1 delta N-ter (27-794) complex 10 x T175 were transfected.

### Protein expression and purification

UBA5, UFC1, UFM1 were expressed and purified as previously described(33). All the fusion constructs (DDRGK1-UFL1, DDRGK1-UFL1ΔN) were expressed in *E. coli* T7 express (New England Biolabs). The transformed cells were grown in 2xYT and induced at 16 **°**C overnight with 0.3 mM isopropyl-β-thio-galactoside (IPTG). For the NMR experiments UFC1 and UBA5 (347-404) transformed cells were grown in standard M9 minimal media supplemented with 15NH_4_Cl and induced at 20 **°**C overnight with 0.3 mM IPTG. The induced cells were harvested by centrifugation at 8000×*g* for 15 min. Pellets were resuspended in lysis buffer (50 mM NaH2PO4 pH 8.0, 500 mM NaCl, 10 mM imidazole, and 5mM β-mercaptoethanol), supplemented with 1 mM phenyl-methyl sulphonyl fluoride (PMSF) and DNase. Cells were disrupted using a microfluidizer (Microfluidics). Lysate was cleared by centrifugation at 20000*g* for 45 min and was subjected to 5 ml His-Trap columns (GE Healthcare). The protein was eluted with a linear imidazole gradient of 15–300 mM in 30 column volumes. Fractions containing the purified protein were pooled and dialyzed overnight at 4°C against dialysis buffer (25 mM NaH_2_PO_4_ pH 8.0, 300 mM NaCl, and 5mM β-mercaptoethanol) in the presence of TEV protease. Cleaved protein was then subjected to a second round of His-Trap column and flow-through containing the cleaved protein was collected. Further purification was done using 16/60 Superdex 75 pg for DDRGK1-UFL1 and DDRGK1-UFL1ΔN, equilibrated in buffer containing Tris-Cl pH 7.5 (20 mM), NaCl (200 mM), and DTT (2 mM). The purified proteins were concentrated and flash-frozen in liquid N2 and stored at −80 °C.

To purify the UFL1 ligase complex from mammalian cells, forty-eight hours after transfection, 293T cells were scraped from T175 flasks, transferred to 50 ml conical tubes, and centrifuged at 1000*g* for 5 min. Cell pellets were resuspended in ice-cold lysis buffer (50 mM Tris-Cl, pH 8, 600 mM NaCl, I mM DTT and 1× protease inhibitor cocktail). The cells are disrupted using microfluidizer and incubated in ice for 30 minutes with 10 μg/ ml avidin, followed by centrifugation at 20,000*g* for 20 min at 4 °C. The supernatant was loaded to 5 ml Strep-tactin superflow HC column. The protein was eluted with 50 mM Tris-Cl, pH 8, 500 mM NaCl, 7.5 mM Desthiobiotin in 10 column volumes. Further purification was done using Superdex 200 pg equilibrated in buffer containing Tris-Cl pH 7.5 (20 mM), NaCl (400 mM), and DTT (1 mM). The purified proteins were concentrated and flash-frozen in liquid N2 and stored at −80 °C.

### *In vitro* ufmylation assay

UBA5 (0.5 μM), His-UFM1 (10 μM), UFC1 (5 μM) and fusion fragments (5 μM each) were mixed together in a buffer containing Hepes (50 mM pH 8.0), NaCl (100 mM) and MgCl_2_ (10 mM). For assay with full length ligase, 0.5 μM of ternary complex and Flag-UFM1 (10 μM) was used. Reactions were initiated by the addition of ATP (5 mM) and were incubated at 30**°**C for 1 hour. The negative control sample was incubated without ATP. After incubation the reactions were stopped by adding 5X SDS-sample buffer containing β-mercaptoethanol. The samples along with the control were then loaded on 10% SDS-PAGE followed by immunoblot with anti-6x His or anti-Flag antibody (Abcam).

### *In vitro* pull down assay

Recombinant purified strep-UFC1 (5 μM) and fusion fragments (5 μM each) were mixed in PBS in total volume of 50 μL for 1 h at RT and subsequently precipitated with Strep-Tactin beads (Iba Lifesciences). The mixtures were washed twice with PBS. The bound proteins were eluted using 7.5 mM desthiobiotin in 50 mM Hepes, pH 8.0 and 300 mM NaCl buffer. Then the samples were analyzed by 4-15% SDS-PAGE followed by Coomassie Brilliant Blue staining.

### Fluorescence-detection size-exclusion chromatography (FSEC)

For the FSEC assay 40 μl of the fragments (10 μM) were injected at flow rate of 0.4 ml min^−1^ to a Superdex 75 Increase 10/300 GL column (GE Healthcare) equilibrated with buffer containing 20 mM Tris-Cl pH 7.5, 200 mM NaCl with 1 mM DTT. Fluorescence was detected using the RF-20A fluorescence detector for HPLC (Shimadzu, Japan) (for Trp, excitation: 285 nm, emission: 330 nm).

### Crystallography

Crystals of DDRGK1-UFL1ΔN were grown at 20°C using the hanging drop vapor diffusion method. Protein was concentrated to 100 mg/ml and crystalized in a solution containing 0.7 M Ammonium tartrate dibasic and 0.1 M Sodium acetate trihydrate, at pH 4.6. The crystals were cryoprotected using a reservoir solution containing 25% glycerol and flash frozen in liquid nitrogen.

Diffraction data for the DDRGK1-UFL1ΔN crystals were collected on beamline ESRF ID30A-3 at 100°K. Data was processed using Dials (34) and scaled using Aimless(35). The putative space group was determined to be hexagonal P6_2_ or its enantiomorph. The structure was solved by molecular replacement (MR) with MOLREP (36) using the AlphaFold2 model of the UFL1-DDRGK1 heterodimer (**Figure 1a**). The translation function confirmed the space group to be P6_4_. The asymmetric unit contains two chains of the chimeric protein. The MR model was refined in REFMAC5 (37) and BUSTER. The electron density was subject to density modification with NCS averaging using Parrot (38, 39). The model was further refined using REFMAC5 with input density modification phases (40). The model was rebuilt using COOT (39) and ISOLDE (41) implemented in ChimeraX (28). Details of the quality of the refined model are presented in **Supplementary Table 1**.

### NMR spectroscopy

All NMR experiments were carried out at 25°C on a 23.5T (1000 MHz) Bruker spectrometer equipped with triple resonance (x,y,z) gradient cryoprobe. The experiments were processed with NMRPipe^58^ and analyzed with NMRFAM-SPARKY^59^. The interaction of UFC1 with DDRGK1-UFL1 fragments was monitored by 2D ^1^H–^15^N HSQC experiments with the assignments for UFC1 transferred from the BMRB [entry 6546 (25)]. DDRGK1-UFL1 (100-400 μM) or DDRGK1-UFL1ΔN (400 μM) were titrated into 200 μM of ^15^N-labeled UFC1 in 20 mM TRIS pH 7.6. For competition experiments, we used ^15^N-labeled UBA5 (residues 347-404; 150 μM), UFC1 (300 μM) and DDRGK1-UFL1 (700 μM).

### Isothermal titration calorimetric experiments (ITC)

ITC experiments were performed using PEAQ-ITC (Malvern Instruments, Malvern) at 25°C. The binding experiments between UFC1 and fusion constructs were performed in buffer containing 20 mM Tris-Cl, 150 mM NaCl, 1 mM DTT, at pH 7.5. Fusion constructs (0.8 mM) were titrated into 80 μM UFC1 *via* 19 individual injections of 2 μl volume each, following the first injection (0.4 μl) that was disregarded. An interval of 120 s was allowed between each injection, and the stirring speed was set at 750 rpm to achieve complete thermodynamic equilibration. The experiment data were analysed with MicroCal PEAQ-ITC analysis software, and “1 set of sites” models were used for data fitting.

## Acknowledgment

This work was supported, in whole or in part, by the Israel Science Foundation, founded by the Israel Academy of Science and Humanities (grant number 491/2021 to R.W. and grant number 301/2021 to O.S.-F.), and by the Israel Cancer Research Fund (award ID 21-113-PG to R.W.). J.K.V. is supported by a Marie Sklodowska-Curie European Training Network Grant #860517 (Ubimotif).

