## Supplementary material for "Structural study of UFL1-UFC1 interaction uncovers the importance of UFL1 N-terminal helix for ufmylation"

### Supplementary Tables

##### Supplementary Table 1. Data collection and refinement statistics

| | |
|---|---|
| Beamline | ESRF ID30A-3 / MASSIF-3 |
| Wavelength (Å) | 0.9677 |
| Space group | $P6_4$ |
| Unit Cell a, b, c (Å), α, β, γ (°) | 145.6, 145.6, 83.2, 90, 90, 120 |
| Resolution range (Å)[a] | 63.03 – 3.07 (3.28 - 3.07) |
| Total reflections [a] | 111,204 (20,857) |
| Unique reflections [a] | 18,967 (3,414) |
| Completeness (%)[a] | 99.8 (99.2) |
| Multiplicity [a] | 5.9 (6.1) |
| $R_{meas}$ (%) [a,b] | 11.5 (179.1) |
| $\langle I\rangle/\langle\sigma(I)\rangle$ [a] | 8.7 (0.5) |
| $CC_{1/2}$ [a,c] | 0.997 (0.446) |
| Wilson B-factor [d] (Å$^2$) | 116.4 |
| $R_{work}$ | 0.231 |
| $R_{free}$ | 0.272 |
| No. of protein monomers in a.u. | 2 |
| Number of atoms | |
| Macromolecules | 4073 |
| Number of protein residues | 511 |
| RMS bond lengths (Å) | 0.015 |
| RMS bond angles (°) | 1.74 |
| Ramachandran favored (%)[f] | 95.7 |
| Ramachandran allowed (%) | 4.3 |
| Ramachandran outliers (%)[f] | 0.0 |
| Clashscore [e] | 4.4 |
| Average B-factor protein (Å$^2$) | 124.1 |
| RCBS PDB code | 8B9X |

[a]Values for the highest resolution shell are given in parentheses

[b] $R_{meas} = \Sigma_h[m/(m - 1)]^{1/2} \Sigma|_i I_{h,i} - \langle I_h\rangle| / \Sigma_h \Sigma_i I_{h,i}$

[c] $CC_{1/2}$ is defined in (Karplus & Diederichs, 2012).

[d] Wilson B-factor was estimated by SFCHECK. (Vaguine et al., 1999).

[e] The Ramachandran statistics and clashscore statistics were calculated using MOLPROBITY. (Williams et al., 2018).


Karplus, P.A., Diederichs, K. (2012) Linking crystallographic model and data quality. Science, 336, 1030-1033.

Tickle, I.J., Flensburg, C., Keller, P., Paciorek, W., Sharff, A., Vonrhein, C., and Bricogne, G. (2018). STARANISO (Global Phasing Ltd.).

Vaguine, A.A., Richelle, J., and Wodak, S.J. (1999). SFCHECK: a unified set of procedures for evaluating the quality of macromolecular structure-factor data and their agreement with the atomic model. Acta Crystallogr. D Biol. Crystallogr. 55, 191–205.

Williams, C.J., Headd, J.J., Moriarty, N.W., Prisant, M.G., Videau, L.L., Deis, L.N., Verma, V., Keedy, D.A., Hintze, B.J., Chen, V.B., Jain, S., Lewis, S.M., Arendall, W.B. 3rd, Snoeyink, J., Adams. P.D., Lovell, S.C., Richardson, J.S., Richardson, D.C. (2018). MolProbity: More and better reference data for improved all-atom structure validation. Protein Sci. 27, 293-315.

**Supplementary Table 2. Interface hotspots at the UFL1-UFC1 interface.** List of residues that contribute significantly to binding, as identified by computational alanine scanning on the model of the UFC1-UFL1 N-terminal helix interface. Bold highlights the residues with a predicted ΔΔG of >1.5 kcal/mol, italics the weaker but still useful residues with ΔΔG >1.0 kcal/mol.

| **UFL1** | | | | **UFC1** | | | |
|---|---|---|---|---|---|---|---|
| Position | Amino acid | ΔΔG(complex) (kcal/mol) | ΔG(partner) (kcal/mol) | Position | Amino acid | ΔΔG(complex) (kcal/mol) | ΔG(partner) (kcal/mol) |
| **5** | **W** | **2.48** | -0.5 | *29* | *V* | *1.23* | 0.36 |
| **8** | **I** | **2.48** | 0.35 | *33* | *K* | *1.07* | -0.1 |
| **11** | **L** | **1.55** | 0.58 | **36** | **Y** | **3.3** | 1.44 |
| **15** | **F** | **4.28** | -0.37 | **40** | **I** | **1.72** | 0.28 |
| *19* | *Q* | *1.17* | 0.5 | *47* | *K* | *1.42* | *-0.13* |
| | | | | **55** | **R** | **1.98** | -0.1 |

T. Kortemme, D. E. Kim, D. Baker (2004). Computational alanine scanning of protein-protein interfaces. Sci. STKE 2004, pl2 (2004).

**Supplementary Table 3. Thermodynamic parameters of binding experiments measured by isothermal titration calorimetry**

| | | |
|---|---|---|
| **Experiment** | ufc1_UFL1 | ufc1_ΔN-UFL1 |
| **Temperature (°C)** | 25.1 | 25 |
| **Bin** | Binding | No binding |
| **[Syr] (M)** | 8.00E-04 | 8.00E-04 |
| **[Cell] (M)** | 8.00E-05 | 8.00E-05 |
| **Control Type** | Fitted Offset | Fitted Offset |
| **N (sites)** | 0.932 | N/A |
| **KD (M)** | 2.29E-06 | N/A |
| **ΔH (kJ/mol)** | -52 | N/A |
| **ΔG (kJ/mol)** | -32.2 | N/A |
| **-TΔS (kJ/mol)** | 19.7 | N/A |

| | | |
|---|---|---|
| **Offset (kJ/mol)** | -0.27 | N/A |
| **Red. Chi-Sqr. (kJ/mol)²** | 0.106 | N/A |

### Supplementary Figures

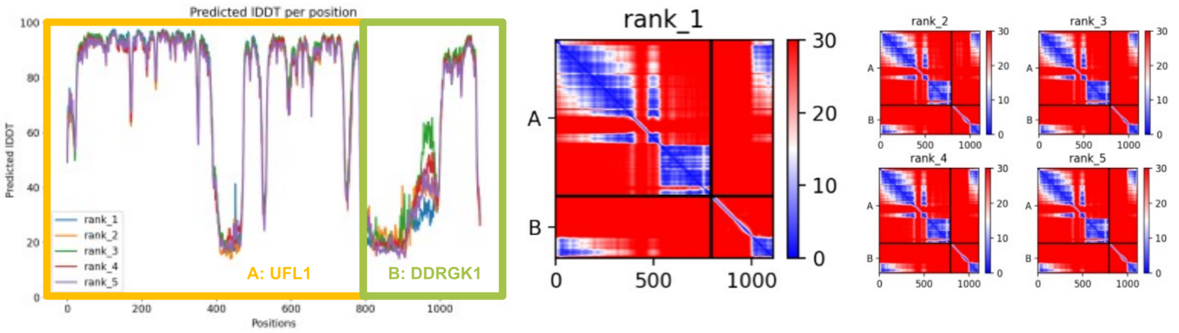


**Supplementary Figure 1. Model confidence of the UFL1-DDRGK1 AlphaFold2 complex prediction. a** Model of the complex formed by separate chains, UFL1 and DDRGK1; **b** Model of the **DDRGK1-UFL1** fusion construct as fusion. Left: Predicted local distance difference test (lDDT); Right: predicted Align Error (pAE) plots of model 1, and models 2-5. Note the high confidence of the fusion construct, including the interface between the two partners (blue areas). For visualization purposes, only rank_1 was used in **Figure 1**. Accompanying **Figure 1a,c**.

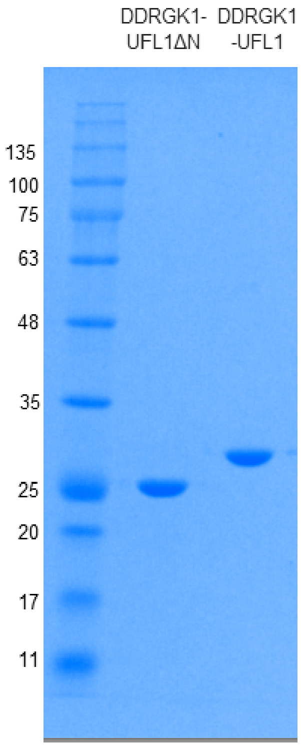


**Supplementary Figure 2.** SDS-PAGE showing the purity of the indicated proteins

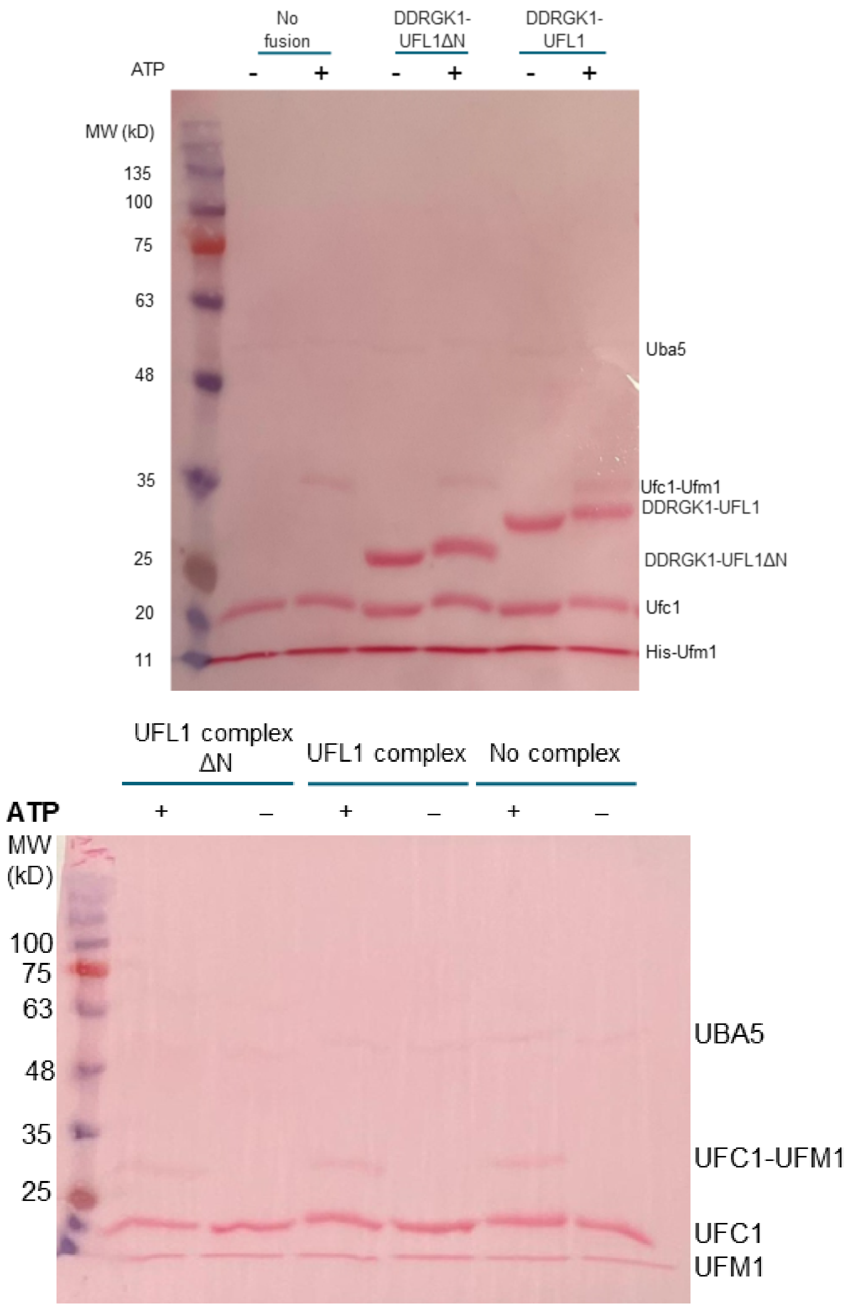


**Supplementary Figure 3.** Loading control of *in vitro* ufmylation assay. **top:** fusion constructs, **bottom:** ternary complex. Following the *in vitro* ufmylation assay the reaction samples were loaded on SDS-PAGE and transferred to a PVDF membrane for Western blot analysis. The membrane was stained with Ponceau S to show equal loading of all proteins in each reaction. Accompanying **Figures 1f&g**.

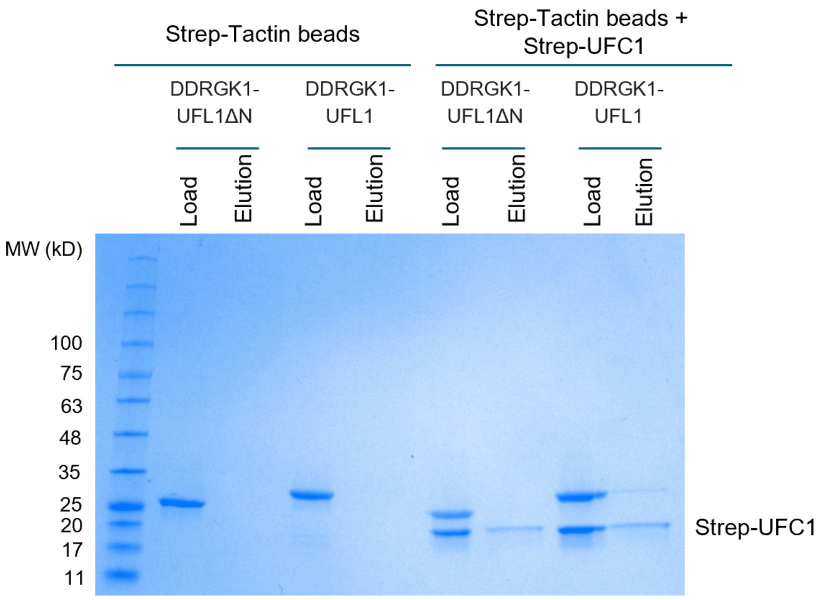


**Supplementary Figure 4.** Coomassie stain gel shows that UFC1 binds to DDRGK1-UFL1 but not DDRGK1-UFL1ΔN, demonstrating that this interaction depends on the presence of the N-terminal helix of UFL1.

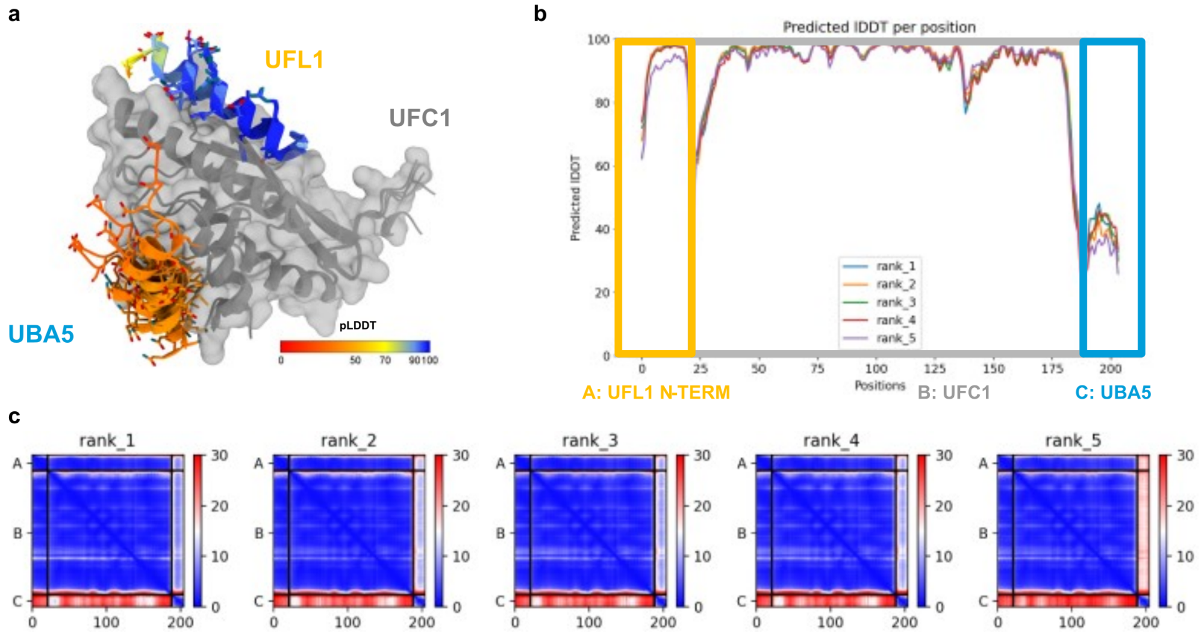


**Supplementary Figure 5. E3 UFL1 and E1 UBA5 compete for the same binding site on E2 UFC1. a** Computational competition assay using AlphaFold2 modeling of the structure in the presence of both helices (N-terminal helix of UFL1 and C-terminal helix of UBA5) identifies UFL1 as the stronger binder, forcing UBA5 to another, incorrect location. Note the high pLDDT for the UFL1 helix (dark blue shades, >90), compared to the corresponding pLDDT values for UBA5 (orange shades, ~50). **b-c** High confidence prediction for UFL1-interaction, compared to UBA5 interaction. **b** pLDDT plot, and **c** pAE plots for the 5 models. Accompanying **Figure 4**.
